# *hURAT1* Transgenic Mouse Model for Evaluating Targeted Urate-Lowering Agents

**DOI:** 10.1101/2023.11.15.567192

**Authors:** Gejing De, Weiyan Cai, Miyi Yang, Qinghe Zhao, Guohua Yi, Peihui Lin, Apeng Chen

**Affiliations:** Institute of Chinese Materia Medica, China Academy of Chinese Medical Sciences, Dongcheng District, Beijing, 100700, China; Department of Medicine, The University of Texas at Tyler School of Medicine, Tyler, TX75708, USA; Center for Biomedical Research, The University of Texas Health Science Center at Tyler, Tyler, TX 75708, USA; Department of Cellular and Molecular Biology, The University of Texas Health Science Center at Tyler, Tyler, TX 75708, USA; Department of Internal Medicine, Division of Nephrology, Davis Heart and Lung Research Institute, The Ohio State University Wexner Medical Center, Columbus, OH 43210, USA; State Key Laboratory for Animal Disease Control and Prevention, Lanzhou Veterinary Research Institute, Chinese Academy of Agricultural Sciences, Lanzhou, 730046, China

**Keywords:** URAT1, *SLC22A12*, hyperuricemia, uric acid, gout, transporter

## Abstract

**Objective:** Urate transporter 1 (URAT1), a well-established urate-lowering therapeutic target for hyperuricemia and gout treatment, expresses in the kidney proximal tubule and is responsible for uric acid (UA) reabsorption. However, non-primate animal models currently used in pharmacological studies failed to evaluate URAT1 inhibitor’s effectiveness because their URAT1 has a very low UA affinity compared to human URAT1, resulting in a lag in targeting drug screening and novel therapy development for gout treatment. We established a human URAT1 (*hURAT1*) transgenic knock-in (KI) mouse model to assess uricosuric agents’ effectiveness and characterize URAT1-caused pathogenesis.

**Methods:** We generated *hURAT1* transgenic mice using CRISPR/Cas9 knock-in technique. *mUrat1* knockout was achieved by replacing exon 1 coding sequence with a human *SLC22A12* CDS-pA cassette. Based on the above transgenic mice, a hyperuricemia model was further established by hypoxanthine administration.

**Results:** The *hURAT1*-KI mice successfully expressed hURAT1 protein to the apical side of the kidney proximal tubule epithelium, where a native human URAT1 kidney localization in human body. Upon hypoxanthine challenge, the blood UA level was elevated in *hURAT1-*KI mice, exhibiting an approximately 37% increase compared to *wild-type (WT)* mice. The elevated blood UA level could be alleviated by hURAT1 inhibitor benzbromarone treatment in the *hURAT1*-KI mice whereas no response was observed in *WT* littermates. Therefore, *hURAT1* transgenic mice responded well to inhibitors and can be used to evaluate the therapeutic effects.

**Conclusion:** The *hURAT1*-KI hyperuricaemia mouse model would be valuable for preclinical evaluation of urate-lowering agents toward gout treatment and studying UA metabolic complexities in humans.

## Introduction

Sustained serum uric acid (UA) elevation causes hyperuricemia and gout, leading to clinical manifestations such as inflammatory arthritis, urolithiasis, hypertension, cardiovascular diseases, and renal failure (1) . The population-based studies showed that the global prevalence of gout ranges between 1% to 6.8%; hyperuricaemia prevalence is 2∼7 folds higher than gout (2). Long-term urate-lowering therapy is the optimal and curative treatment for gout and hyperuricemia (3). Patients experiencing the first attack of gout flare need to achieve and maintain a target plasma urate within 5 mg/dL or less. Urate-lowering intervention facilitates UA crystal dissolution and prevents the recurrence of gout flares, reducing the risk of tophi, joint damage, or nephrolithiasis (4). Emerging clinical evidence has indicated that achieving a lower serum urate level by therapy will benefit people with gout (3). The probability of being free of gout flare is increased when a lower serum urate is maintained (5). However, limited commercially available medications have not satisfied the global urate-lowering demands, highlighting the urgency of developing such drugs.

Currently, commercial drugs to reduce blood urate, including xanthine oxidase (XO) inhibitors and urate transporter inhibitors representing respective categories, are based on distinct mechanisms: XO inhibitors reduce uric acid production by blocking xanthine to uric acid conversion; Transporter inhibitors increase body uric acid elimination by blocking uric acid recycling. URAT1, encoded by *SLC22A12* gene, is a urate-anion exchanger that regulates blood urate homeostasis by regulating most kidney urate reabsorption from the original urine in the late part of the proximal tubule (6). URAT1 contributes to more than 50% of body uric acid maintenance; hence, it is the most important target for developing urate-lowering drugs. Clinical evidence showed that individuals carrying two nonfunctional *hURAT1* alleles developed an 88% decrease of blood urate level (7). Moreover, 90% of hyperuricemia patients were caused by the exertion impediment rather than the urate overproduction (8). Hence, URAT1 inhibitors as uricosuric treatments showed a better performance than XO blockers for those people with gout (9). Nonetheless, only a few URAT1-targeted drugs have been developed and proven effective in clinical settings in the past decades. Benzbromarone and lesinurad are two URAT1 inhibitors that have been approved only in a few countries and regions to treat gout. Although benzbromarone is a potent urate-lowering drug, the USA and European countries prohibit its use due to its severe hepatotoxicity. Lesinurad was withdrawn from the market because of its poor clinical performance in the USA. Therefore, there is an urgent demand to screen and optimize novel URAT1-targeted drug candidates with improved bioavailability and less toxicity.

UA is a powerful antioxidant and neuroprotector (10). Higher UA levels create more congenial physiology environments for enhanced intelligence, maintaining blood pressure, alleviating oxidative stress, and being fit for fructose consumption, which is considered an evolutionary advantage in modern apes. Nevertheless, when the serum UA concentration exceeds the saturation threshold (> 7.0 mg/dL), it forms monosodium urate crystal and deposits in the joint, kidney, and multiple organs, causing harmful or deadly clinical consequences. The fine-tuning URAT1 occurring in primate species counteracts severe consequences of functional uricase loss and achieves plasma urate homeostasis (11). Rodent URAT1 orthologs exhibit 14%∼20% binding affinity and 7∼9 folds of transporting capacity for UA as a substrate compared to human URAT1 (11), which is consistent with our observation, thus, it reasonable that m*URAT1* gene knockout mice exhibit no function impairment and serum UA alteration (12, 13). The difference could be caused by secretory mechanisms that determine the varied susceptibility of drugs among species. Human URAT1 affinity for benzbromarone is remarkably higher than rats with IC_50_ 0.13 μM for hURAT1 and IC_50_ 11.9 μM for rURAT1, respectively (14). Our results show that mURAT1 also exhibits a much lower affinity for benzbromarone than hURAT1. As such, *WT* mice and rats rarely respond to hURAT1 inhibitors (i.e., benzbromarone); hence, they are inappropriate for URAT1-related drug screening and evaluation in preclinical studies.

Currently, the New World monkey *Cebus apella* (Brown capuchin) is the only animal model for URAT1 preclinic pharmacology studies (15, 16), which brings a huge financial burden and access difficulty for primary drug screenings and pharmacological activity evaluations. In this study, we developed a *hURAT1* knock-in (*hURAT1*-KI) mouse model to functionally demonstrate the human-like URAT1 reabsorbing capability of uric acid *in vivo*. We further confirmed that the *hURAT1* transgenic mouse is an ideal small animal model for pharmacology development and fundamental research on urate metabolic-associated diseases.

## Results

### Generation and characterization of human *URAT1* knock-in mice

We used CRISPR/Cas9 gene editing to create a *hURAT1*-KI mouse model that mimics human-like uric acid metabolism. The *Slc22a12* gene (NM_009203.3) is located in mouse chromosome 19 GRC m38.p6 (Figure 1A). The *hURAT1* gene coding sequence (CDS) with a simian virus 40 (SV40) polyadenylation signal was subcloned into the vector and microinjected into C57BL/6J genetic background single-cell embryos. *hURAT1* gene inserted and replaced exon 1 coding sequence of mouse *Urat1* (*mUrat1*), resulting in *mUrat1* silence (Figure 1A). Founder animals (F0) having successful insertions were confirmed by PCR screening and genomic DNA sequencing (Figure 1B and C). Five knock-in founders containing the *hURAT1* transgene demonstrated *hURAT1*-specific amplicons. F0 mice were further bred with *WT* mice to establish F1 heterozygous offspring. Southern blot hybridization further validated heterozygosity with correct transgene copy numbers (Figure 1 D). Transcriptional expression of the *hURAT1* and *mUrat1* genes was assessed by RT-PCR (Figure 1E). mRNA of human and murine *URAT1* were expressed in heterozygous (*hURAT1* KI/+) mouse kidney tissue, respectively. In contrast, only *mUrat1* allele was detected in *WT* mice.

**Figure 1.**
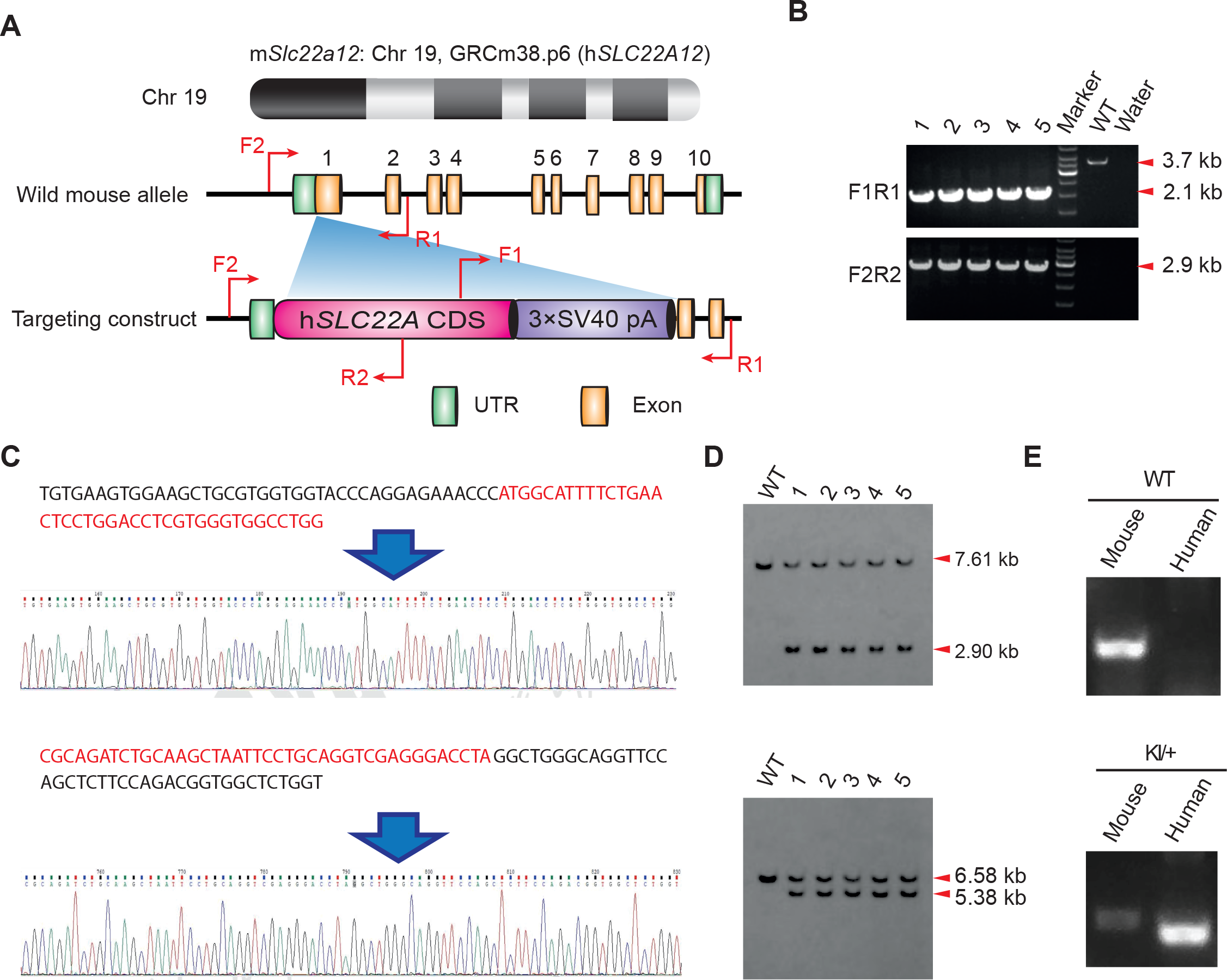
Establish *hURAT1*-KI mouse. (A) Configuration of *hURAT1* cDNA transgene. The *hURAT1* gene was inserted into exon1 of *mUrat1* in chromosome 19 GRC m38.p6. F0 founder animals were identified by PCR screening (B) and DNA sequencing (C), which were bred to *WT* mice to test germline transmission and F1 animal generation. *WT* animals harbor the full-length exon1 of *mUrat1* gene, which extends into the size range of 3.7 kb. Only 2.1 kb size is detected in animals harboring the *hURAT1* transgene. (C) *hURAT1-*specific primer was applied in DNA sequencing in transgenic mice genomic DNA. (D) F1 mice were confirmed by southern blot. The endonucleases *AvrII* and *StuI* were used to identify the correct insertion. Expected fragment sizes for southern blot: 5’Probe-AvrII: WT-7.61 kb, *hURAT1*-2.90 kb; 3’Probe-Bsu36I: *WT*-6.58 kb, *hURAT1*-5.38 kb, respectively. (E) Transcriptional analysis of transgenic and *WT* lines. The positions of the endogenous murine *Urat1, hURAT1*, and *Gapdh* transcripts are indicated. Heterozygous targeted mice expressed both human and mouse *URAT1* genes.

### Phenotype of *hURAT1-*KI mouse

Immunostaining analysis of kidney using an anti-hURAT1 antibody demonstrated the expression of *hURAT1* in homozygous (*hURAT1* KI/KI) and heterozygous (*hURAT1* KI/+) mice. The results showed that human URAT1-specific antibodies do not recognize rat and mouse URAT1 proteins (Figure 2A). Anti-hURAT1 antibody detected highly expressed hURAT1 in *hURAT1* KI/KI) and *hURAT1* KI/+ mice. Expressed hURAT1 is in epithelial cells of proximal tubules in the renal cortex, with tissue distribution similar to that of the human kidney (Figure 2A).

**Figure 2.**
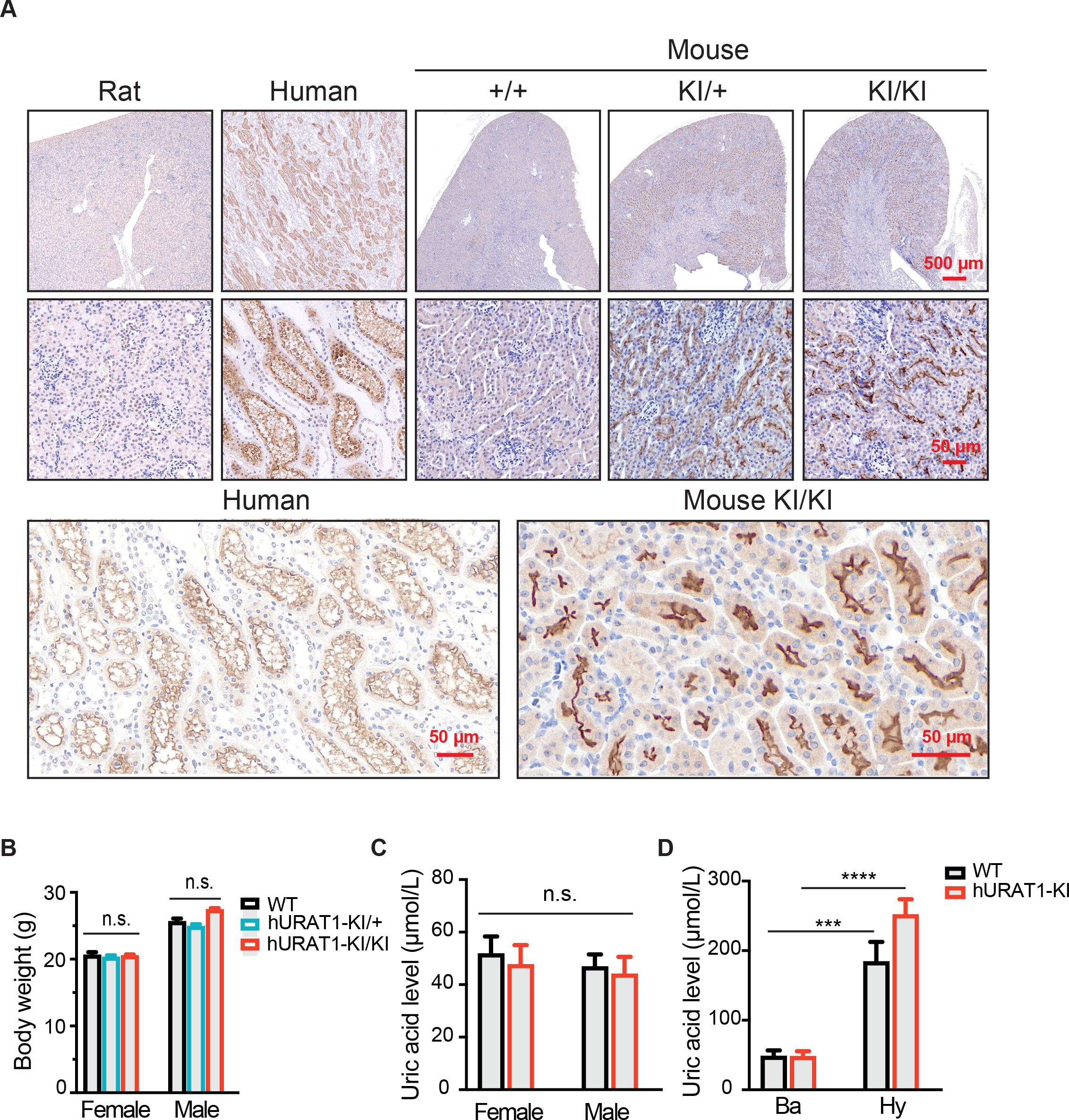
The phenotype of *hURAT1*-KI animals. (A) Immunohistochemistry staining analysis for the hURAT1-expressed in rat, human, and mouse kidney tissues. The primary antibody used here was specifically reacted with hURAT1. The histologic appearance of cortical renal tubules in humans and *hURAT1* mice, was not detected in *WT* mice and rats. Brown color indicates positive staining. (B) Bodyweight of *hURAT1* mice and *WT* littermates. Plasma uric acid concentration of basal metabolic condition (C) and hyperuricemia (D) in *hURAT1* mice and *WT* littermates. Ba, basal; KI, *hURAT* knock-in; Hy, hyperuricemia. n=12 in *WT-ba*, n = 12 in *hURAT1* KI-ba, and n =29 in *hURAT1* KI+Hy mice group, n=20 in *WT* +Hy mice group.

Homozygous hURAT1 transgenic mice are fertile and viable. They displayed a normal life span without apparent phenotypic alterations and only minor increased body weight in male homozygous animals with no significant difference when assessed at age of two months (Figure 2B).

We further analyzed blood UA levels of the *hURAT1*-KI and *WT* mice. UA levels of male and female *hURAT1* mice are comparable to *WT* mice [43.9 µmol/L (male) and 47.5 µmol/L (female) in *hURAT1* mice v.s. 46.7 µmol/L (male) and 51.6 µmol/L (female) in *WT* mice] (Figure 2C). The results showed that *hURAT1*-KI mice maintained similar low-level UA to *WT* mice, which is physiologically reasonable since the functional uricase is in charge of most urate elimination in normal physiological conditions of mice. To further establish an animal model with a blood urate level close to the physiological of the human body, normally ranges 180 - 420 µmol/L, we utilized hypoxanthine to stimulate the mice (Figure 2D). A robust increase in urate level was observed in both *hURAT1-*KI and *WT* mice; interestingly, transgenic mice had a 37% higher value (averaged 251 µmol/L) than the *WT* mice (averaged 183 µmol/L), suggesting that hURAT1 transported more urate from urine to blood compared with mURAT1, subsequently enhancing the blood urate level. The hURAT1 and mURAT1 exhibit 83% similarity in protein sequence, while hURAT1 has roughly ∼5 times higher capability to transport uric acid than mURAT1 (11). Our data confirmed the previous observation that hURAT1 is more effective in regulating physiological UA that range 200 - 400 µmol/L (11).

### *hURAT1*-KI mice for elevating *hURAT1*-targeted drug performance

To validate whether the *hURAT1-*KI mouse could be applied as a tool for evaluating URAT1 inhibitors, we investigated the effectiveness of URAT1 inhibitor benzbromarone, a commercial urate-lowering drug. We established stable HEK293T-hURAT1, HEK293T-mURAT1, and HEK293T-GFP cells for *in vitro* UA transport assay. The result showed that benzbromarone achieved a significant effect at a considerable inhibition rate in hURAT1-expressed cells, 0.18 µM compared to 45 µM in mURAT1-expressed cells (Figure 3A). The data revealed an obvious difference between hURAT1 and mURAT1 in affinity of inhibitor, which is why the *WT* mouse model had no responses to UTAT1 inhibitor treatment. We then tested in our established hypoxanthine-induced hyperuricaemia model to test its sensitivity toward benzbromarone treatment. Benzbromarone showed a decrease of blood UA in *hURAT1-*KI mice (decreased to 63% of non-treatment), while there was no significant change in *WT* mice (92% of non-treatment) (Figure 3B). Our study demonstrated an animal model that could effectively respond to URAT1 blocking, which can be an important preclinical investigation tool to test drug therapeutic effects.

**Figure 3.**
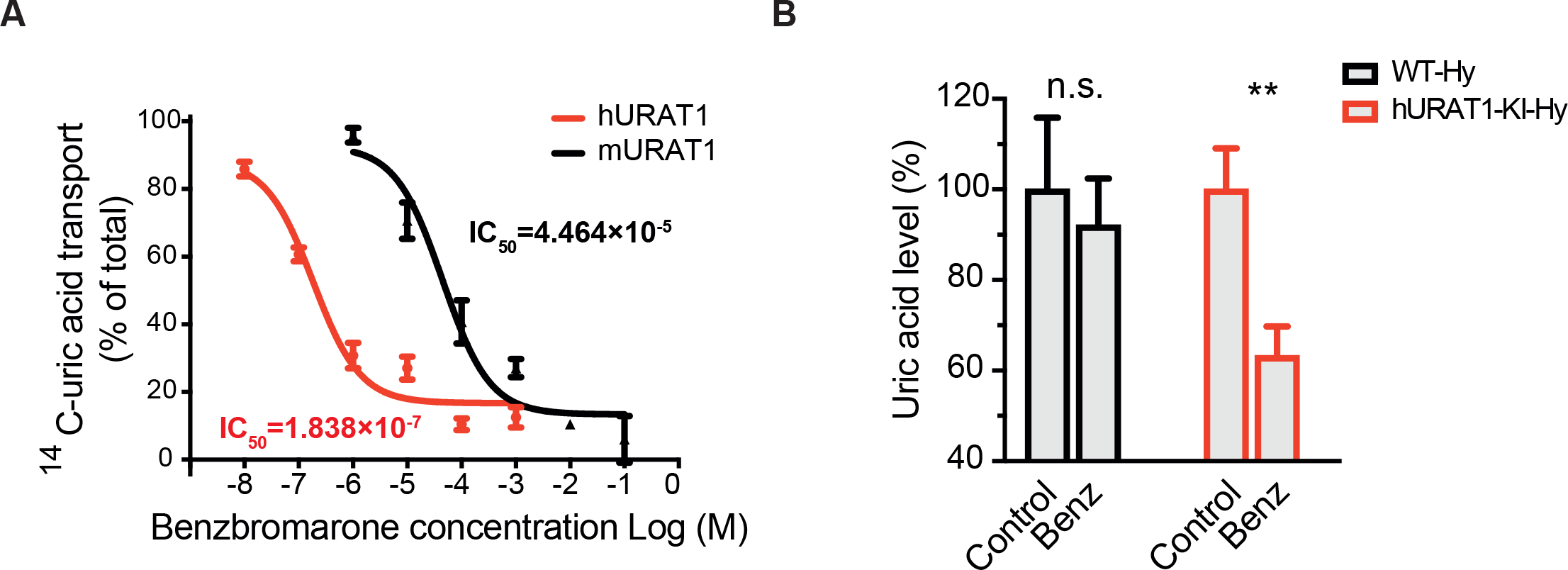
Benzbromarone alleviate uric acid level in hyperuricemia *hURAT1-*KI mice model. (A) Dose responses for benzbromarone to inhibit the transport activity of human (red) and mouse (black) URAT1. Cells expressing URAT1S were treated with ^14^C-uric acid for 20 mins. Results are averaged from two independent experiments, each done in triplicate ± SD. (B) Model function analysis: Benzbromarone lowers plasma UA in *hURAT1-*KI mice. A single dose (480 mg/kg) of benzbromarone, by oral gavage, lowered the plasma UA levels by up to 37%. Results are representative of data generated in two independent experiments and are expressed as mean ± SEM. n=20 in *WT*, n = 15 in *WT*+Benz and n =29 in *hURAT1*-KI mice group, n=29 in *hURAT1*-KI+Benz mice group. Statistical significance was determined with *two-way ANOVA* analysis.

## Discussion

The human UA level normally ranges from 3.5 to 7 mg/dL (equivalent to 200∼400 µmol/L)(17) . Unlike humans, most mammal species’ serum uric acid levels are in the range of 0.5∼1.5 mg/dL (equivalent to 30∼90 µmol/L) range since the uricase enzyme in these species converts uric acid into a more soluble 5-hydroxyisourate as an end-product of purine metabolism, which facilitates final metabolite elimination from the body. 20 million years ago, multiple accumulated nonsense mutations appeared to form a nonfunctional uricase gene during primates’ evolution, making an elevated level of serum urate possible (18). Consequently, humans are susceptible to hyperuricaemia and gout that is caused by abundant urate in blood. The evolutionarily acquired URAT1 appears in great apes (orangutan, gorilla, chimpanzee, and human) in a similar period and species as uricase-deficient events occur, with appropriate substrate binding affinity and low capacity to achieve more serum UA retention and precise control. Primate URAT1 can subtly balance serum urate, hence counteracting blood uric acid elevation caused by uricase functional loss. Inhibiting hURAT1 could suppress UA reabsorption in the kidney proximal tubule, lowering blood UA and preventing monosodium urate crystal formation, ultimately improving the progression of gout and hyperuricemia.

On the other hand, the difference in capacity/affinity of URAT1 among species determines variations in sensitivity to hURAT1-target molecules. The field has suffered from a lack of appropriate animal models expressing human-like URAT1 that could effectively respond to URAT1 blockade, which is the primary reason for the delayed URAT1 investigation and urate-lowering drug development.

In summary, we have developed an animal model that bridges a discrepancy between rodents and primates regarding URAT1 function. Our result demonstrates that benzbromarone treatment could reduce blood UA levels in *hURAT1* mice, increasing urate excretion, while there was no apparent response in *WT* mice. Therefore, *hURAT1-KI* transgenic mouse model represents a convenient and valuable small animal model to evaluate the effectiveness of URAT1-targeting drug candidates.

## Materials and Methods

### *hURAT1*-KI mice generation

Human URAT1 knock-in mice (*hURAT1-*KI) were developed at Cyagen Biosciences Inc. (Suzhou, China) by introducing a donor vector containing *SLC22A12* CDS-P2-3*SV40 pA cassette and Cas9 and gRNA into C57BL/6 embryonic stem cells to generate targeted knock-in offspring. F1 mice were intercrossed to generate homozygous mice. PCR screening primers as following: forward primer (F1): 5’-TCACCATCTACAGCAGCGAGCT-3’ reverse primer (R1): 5’-GGTTCACTCAGTAGAGACCGCCT-3’; forward primer (F2): 5’-CAGGAATCGTACGGACATCTCTAT-3’; reverse primer (R2): 5’-GTTCAAGGTCATCACCAAGGGTC-3’. Genomic DNA sequencing primers were as follows: forward primer 5’-CCAGATTACCACAGAGGGTTCC-3’; reverse primer CTCTGTAAGCTGCCATTGAGGTTG-3’. Genotyping primers were as follows: forward primer-5’-TGGTGCTACTCTGTGGTGCTA-3’; reverse-5’-CAGGAATCGTACGGACATCTCTA-3’.

### Animal studies

All animal studies were conducted following the protocol of the China Academy of Chinese Medical Sciences (No. of approval: 2023B102). Sex and age-matched transgenic and WT littermates were used for experimental analyses.

The baseline plasma urate levels of mice were performed. Tissue samples were obtained for histological analysis. The left kidney of the animal from each group was sectioned longitudinally and subjected to immunohistochemistry.

Mouse hyperuricaemia model: to simulate human body UA levels (ranges 200∼400 µmol/L), we induced a high plasma UA level in mice. Mice were injected with 480 mg/kg hypoxanthine solution (in sterile water). Blood was collected four hours post-injection for analysis. *WT* mice received the same treatment served as controls.

### Cell culture and lentivirus packaging

HEK-293T cells stably expressing human and mouse URAT1 were established. In brief, with the full length of *hURAT1/mURAT1* cDNA were subcloned into plent-EF1a-FH-CMV-GFP vector. HEK-293T cells stably expressing hURAT1 (as HEK293T-hURAT1) or mURAT1 (as HEK293T-mURAT1) were obtained by viral infection of HEK-293T cells with packaged lentiviral transgenes as above-mentioned. HEK-293T cells transduced with packaged lentivirus with plent-EF1a-FH-CMV-GFP vector were used as a control (HEK293T-GFP mock cells). The viral-containing medium was changed to puromycin selection media 24 h after infection. Survived stable cell lines were grown in a humidified incubator at 37 oC and 5% CO2 using a minimum essential medium containing 10% fetal bovine serum and 2 µg/mL puromycin. Urate uptake experiments were performed by following the protocol as previously described (19).

### Blood sample collection and analysis

Whole blood was collected from homozygous *hURAT1-*KI and *WT* mice into commercially available EDTA-treated tubes. Plasma was harvested by centrifugation of the whole blood for 10 min at 2500 rpm. Following centrifugation, the upper layer plasma was transferred into aliquots and stored at -20 °C for future analyses. Plasma urate levels were measured using the Uric Acid Kit (C012-2, Nanjing Jiancheng Bioengineering Institute, Nanjing, China) by following the manufacturer’s protocol. For each set of measurements, a standard curve was generated.

### Immunohistochemistry and Imaging

Mice were euthanized for tissue collection. Kidneys of animals from each group were fixed in 10% neutral buffered formalin, paraffin-embedded, sectioned and subjected to histological (hematoxylin-eosin (H&E) stain) and Immuno-histochemical (IHC) analyses. Kidneys were stained with a hURAT1-specirfic antibody (Sino Biological Inc. Beijing. China) for expression analysis. Sample images were taken using a Pannoramic Scan instrument (3DHISTECH Kft.; Budapest, Hungary).

## Statistical analysis

Statistical analyses were performed using GraphPad Prism version 5.0 (GraphPad Software, San Diego, CA, USA). The two-way ANOVA was used to compare multiple groups. The figure asterisk indicates the statistical significance (*, *p-*value <0.05; **, *p-* value <0.01; ***, *p-*value <0.001).

## Abbreviations

UA: uric acid
URAT1: urate transporter 1
RT-PCR: reverse transcription-PCR
i.p. injection: intraperitoneal injection
ns: not significant
Ctrl: control

## Author Contributions

G De conceived the project and designed the research; A Chen coordinated resources; G De and W Cai performed research; G De and A Chen wrote the manuscript; G Yi and P Lin revised the manuscript.

## Acknowledgments

This work was supported by the National Natural Science Foundation of China (Program, Grant NO. 32273019) grant and the Natural Science Foundation of Gansu Province (grant no.22JR5RA028) grant awarded to A Chen, The Major Science and Technology Project of Gansu Province (22ZD6NA001), grants from the National Natural Science Foundation of China (Grant NO. ZXKT23002) awarded to G De.

